# First fully-automated AI/ML virtual screening cascade implemented at a drug discovery centre in Africa

**DOI:** 10.1101/2022.12.13.520154

**Authors:** Gemma Turon, Jason Hlozek, John G. Woodland, Kelly Chibale, Miquel Duran-Frigola

## Abstract

We present ZairaChem, an artificial intelligence (AI)- and machine learning (ML)-based tool to train small-molecule activity prediction models. ZairaChem is fully automated, requires low computational resources and works across a broad spectrum of datasets, ranging from whole-cell growth inhibition assays to drug metabolism properties. The tool has been implemented end-to-end at the Holistic Drug Discovery and Development (H3D) Centre, the leading integrated drug discovery unit in Africa, at which no prior AI/ML capabilities were available. We have exploited in-house data collected from over a decade of drug discovery research in malaria and tuberculosis and built models to predict the outcomes of 15 key checkpoint assays. We subsequently deployed these models as a virtual screening cascade at an organisational scale to increase the hit rate of current experimental assays. We show how computational profiling of compounds, prior to synthesis and experimental testing, can increase the rate of progression by up to 40%. Moreover, we demonstrate that the approach can be applied to prioritise small molecules within a chemical series and to assess the likelihood of success of novel chemotypes, promoting efficient usage of limited experimental resources. This project is part of a first-of-its-kind collaboration between the H3D Centre, a research centre operating in a low-resource setting, and the Ersilia Open Source Initiative, a young tech non-profit devoted to building data science capacity in the Global South.

## Introduction

The cost of bringing new medicines from the bench to the bedside has risen steadily since the 1970s (DiMasi, Grabowski, and Hansen 2016). Recent estimates suggest a median cost of $1.1 billion per drug (Wouters, McKee, and Luyten 2020) with research and development taking an average of ten years (Brown et al. 2022). To tighten its research pipelines and avoid costly failures, the drug discovery industry has turned to artificial intelligence (AI) and machine learning (ML) to accelerate research timelines and reduce attrition rates. The application of AI/ML aims to transform drug discovery from a slow, sequential, high-risk process to a fast, finely-tuned and integrated pipeline that expedites the delivery of novel clinical candidates with reduced risk of failure (Vamathevan et al. 2019). Notably, investment in AI/ML-based drug discovery by pharmaceutical companies and biotech startups has soared in the last five years (Kirkpatrick 2022), bringing over 150 small molecules to discovery pipelines (Jayatunga et al. 2022).

The promise of AI/ML for biomedicine extends to the field of infectious diseases, which are currently underrepresented in drug discovery portfolios (WHO 2022). Infectious diseases predominantly afflict lower-to-middle-income countries (LMICs), most of which are situated in the Global South. For example, Africa carries over 95% of the 240 million annual global cases of malaria (World Malaria Report 2021) and 25% of global deaths from tuberculosis (TB) (Jeremiah et al. 2022). Historically, the pharmaceutical industry’s efforts to tackle these challenges have principally occurred in the Global North; consequently, African drug discovery efforts have largely been dependent on international funding agencies with programmes driven from abroad. AI/ML methods offer an opportunity to revitalise and expedite drug discovery projects conducted in those countries that are most affected by these scourges; however, a lack of data science expertise and limited access to computational resources hinder the uptake of AI/ML at research institutions and universities in LMICs (Alami et al. 2020). It is anticipated that lowering these barriers may lead to important scientific contributions from those countries that disproportionately suffer from the bulk of infectious diseases, a milestone towards their eradication.

Since its launch in 2010, the Holistic Drug Discovery and Development (H3D) Centre at the University of Cape Town in South Africa has made significant advances in innovative drug discovery projects and infrastructure development, intimately aligned with capacity strengthening across the African continent (Winks et al. 2022). This includes the discovery of the first ever small-molecule clinical candidate, for any disease, researched on African soil by an international team led by an African drug discovery centre (Nordling 2013) which, subsequently, reached Phase II human trials in African malaria patients. To advance its mission of discovering and developing novel, life-saving medicines for infectious diseases that predominantly affect African populations, H3D works closely with the Ersilia Open Source Initiative (EOSI), a non-profit organisation aimed at disseminating AI/ML methodologies applied to urgent biomedical needs in LMICs.

Here we describe ZairaChem^1^, an automated pipeline for AI/ML modelling, designed for fast and easy implementation in low-resource settings. We demonstrate the application of ZairaChem to key assays in the antimalarial and antitubercular drug discovery programs conducted at H3D. The AI/ML models are arranged in the form of a virtual screening cascade that mirrors the progression of compounds in a real-world experimental setting. Virtual screening is a critical tool in drug discovery, allowing for the identification of new hits at a fraction of the time and cost associated with experimental assays. Large chemical libraries can be filtered through this computational cascade to prioritise compounds for testing, resulting in a significant reduction in experimental attrition rates. The developed AI/ML assets include models corresponding to whole-cell phenotypic screening assays against *Plasmodium falciparum* (*Pf*) and *Mycobacterium tuberculosis* (*Mtb*), the causative agents of malaria and tuberculosis, respectively. The cascade also supports later phases of the drug discovery pipeline aimed at de-risking hits with respect to liabilities such as microsomal clearance and cytotoxicity.

This is the first comprehensive AI/ML-based virtual screening cascade that, to our knowledge, has been brought to production in a drug discovery setting on the African continent. ZairaChem can run on conventional computers and is fully automated, requiring limited data science expertise and allowing for periodic model updates with new data. We believe that this virtual screening cascade, based on in-house data and free open-source software, has the potential to set the basis for sustainable, affordable and scalable drug discovery initiatives in the Global South.

## Results

### Available data and screening cascades at H3D

We have modelled two virtual drug discovery cascades focused on identifying novel antimalarial and antituberculosis compounds. Both cascades start with whole-cell activity assays and progress with common absorption, distribution, metabolism, excretion and toxicity (ADMET) profiling (Fig. 2a). We have selected assays representative of each step of the experimental drug screening cascade and for which H3D had sufficient available in-house data (i.e. at least 100 molecules). Key assays for compound progression in the cascade but for which only a small subset of data was available (e.g. human cytochrome P450 (CYP) inhibition and hERG blockade) have been developed using publicly available data (see Materials and Methods). Drug metabolism and pharmacokinetics (DMPK) assays, characteristic of a lead optimisation phase in the drug discovery pipeline, have not been included in the virtual screening cascade at this stage of implementation. Extended Data Table 1 contains a summary of each assay and the number of compounds available. Expert knowledge was used to establish a measurement cut-off to obtain binary outcomes (1/0; active/inactive, soluble/not soluble, etc.) for each of the assays.

**Figure 1.**
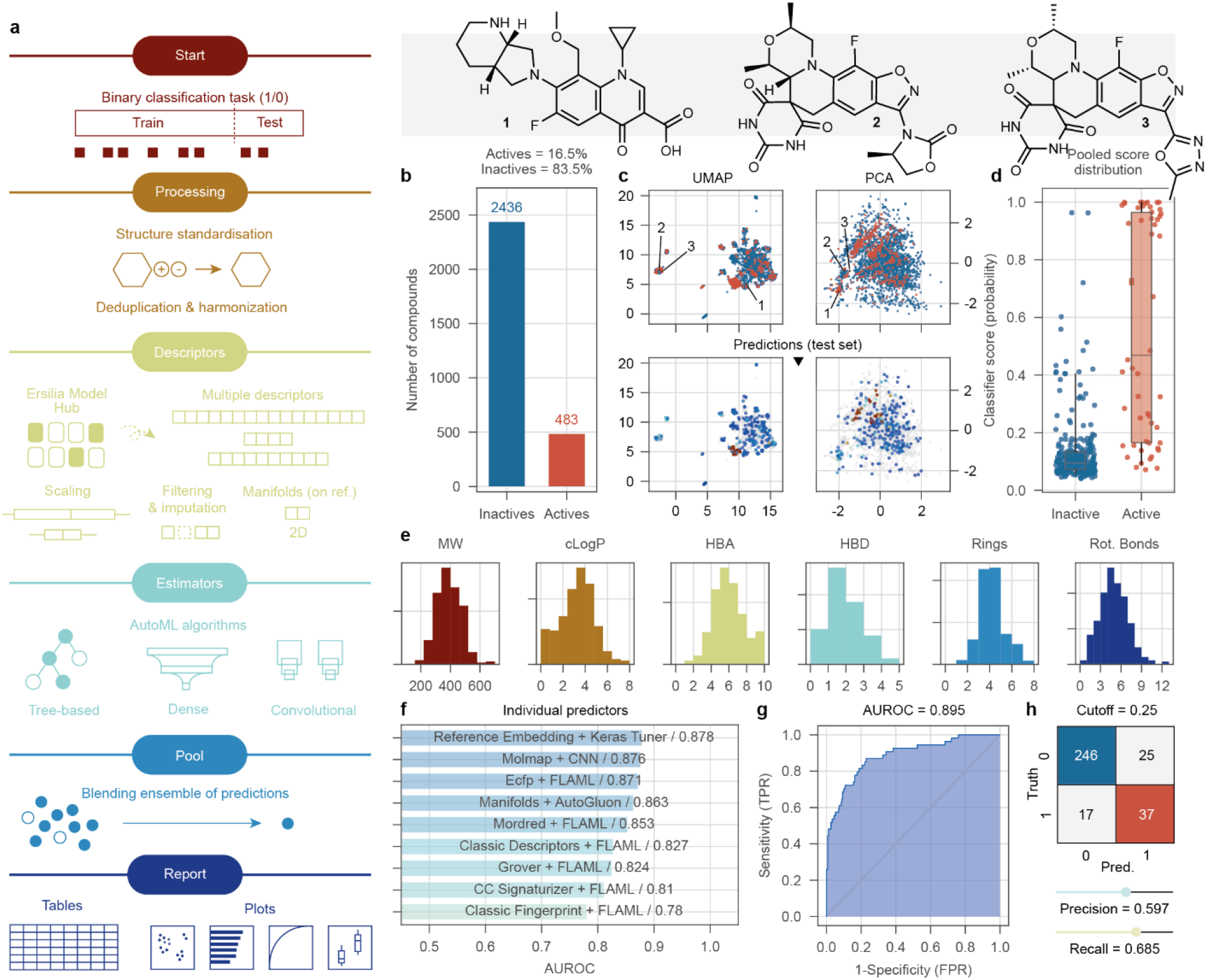
The ZairaChem pipeline. (**a**) Scheme of the AutoML methodology, consisting of data processing, descriptor calculation, training of models, assembling (pooling) of results and reporting. (**b**) Number of active and inactive compounds in the *Mtb* MIC_90_ assay (training set). (**c**) Uniform manifold approximation and projection (UMAP) and principal component analysis (PCA) projections of the chemical space in the *Mtb* MIC_90_ assay. Structurally different (1 vs 2/3) and similar (2 vs 3) compounds are depicted. Red indicates active compounds; blue indicates inactive compounds. (**d**) Model scores (probability of “1”) assigned to the true active (red) and inactive (blue) compounds in the test set (10% of the total available data). (**e**) Distribution of common chemical properties of the compounds, namely molecular weight (MW), calculated logP (cLogP), number of hydrogen bond acceptors (HBA), number of hydrogen bond donors (HBD), number of rings (Rings) and number of rotatable bonds (Rot. Bonds). (**f**) AUROC scores of the individual ZairaChem predictors. (**g**) ROC curve of the final ensemble model. (**h**) Confusion matrix showing true positives (red), true negatives (blue), false positives and false negatives in the test set.

**Figure 2.**
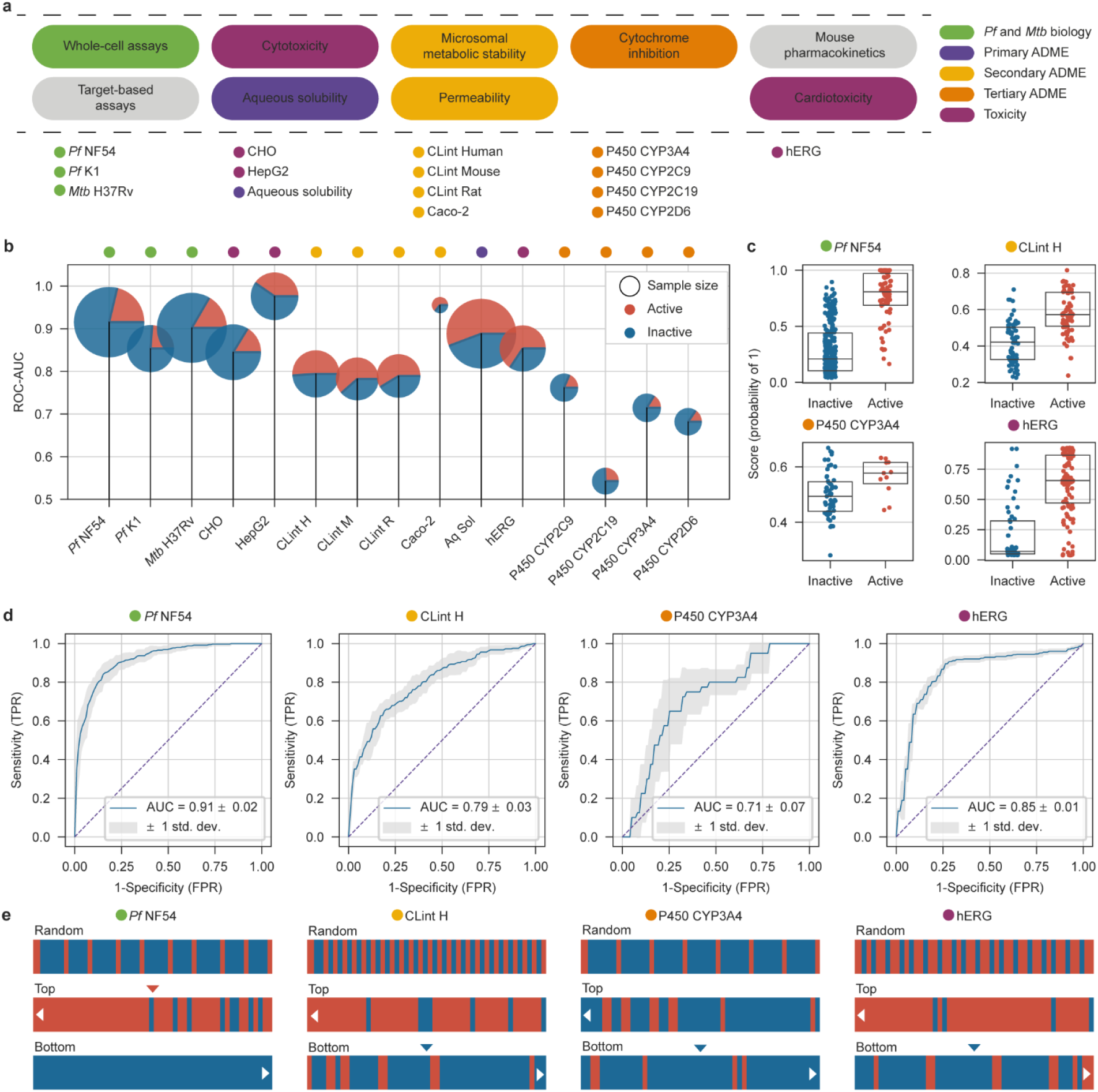
ZairaChem implementation of a virtual screening cascade. (**a**) Summary of assays most frequently used in drug discovery programmes at the H3D Centre, progressing from left to right. (**b**) AUROC for the 10 AI/ML models developed with internal data (test sets: 10% of H3D data), the four CYP models developed with ZairaChem using external data, and the CardioToxNet model from the literature (test sets: 100% of H3D data). Dataset sample counts are represented by circle size with the corresponding proportion of active (red) and inactive (blue) compounds. (**c**) Classification scores of individual compounds for representative assays of different stages of the screening cascade. (**d**) Correspondingly, ROC curves resulting from a five-fold cross-validation, with blue lines depicting the mean AUROC. (**e**) Comparison of hit rates for randomly selected molecules (first row) vs molecules ranked according to the model score (probability of “1”) for selected assays. The top 50 (second row) and bottom 50 (third row) molecules are depicted, showing a hit enrichment of true active compounds (red) in the highest-ranked positions and an enrichment of true inactive compounds (blue) in the lowest-ranked positions. Red and blue arrows, respectively, represent the desired experimental outcome for molecule progression in the cascade (e.g. active *Pf* NF54 molecules and non-active molecules in the hERG assay are desired biochemical properties).

### Automated AI/ML modelling with ZairaChem

Using known binary (1/0) outcomes, we trained robust AI/ML classification models able to predict the probability of “1” (typically an “active” assay outcome) of new compounds, given only their chemical structures represented as a SMILES string. To streamline model training and facilitate adoption of the AI/ML models at H3D, we developed an automated AI/ML tool named ZairaChem. Details of the ZairaChem pipeline are provided in the Materials and Methods section, and a scheme of the methodology is presented in Figure 1a. In brief, molecules are represented numerically using a combination of distinct descriptors, including physicochemical parameters (Mordred (Moriwaki et al. 2018)), 2D structural fingerprints (ECFP (Rogers and Hahn 2010)), inferred bioactivity profiles (Chemical Checker (Duran-Frigola et al. 2020)), and graph-based embeddings (GROVER (Rong et al. 2020)). The rationale is that combining multiple descriptors will enhance applicability over a broad range of tasks, ranging from aqueous solubility prediction to phenotypic outcomes. Subsequently, a battery of AI/ML algorithms is applied using modern automated learning (AutoML) techniques aimed at yielding accurate models without the need for human intervention (i.e. algorithm choice, hyperparameter tuning, etc.). Automation is key to ensure continuous integration and continuous deployment of the AI/ML assets in an environment like H3D, where data science capacity is limited. The AutoML frameworks FLAML (Wang et al. 2019), AutoGluon (Erickson et al. 2020) and Keras Tuner (O’Malley et al 2019) were incorporated, covering mostly tree-based methods (Random Forest, XGBoost, etc.) and neural network architectures. We benchmarked the ZairaChem pipeline in the Therapeutics Data Commons ADMET binary classification tasks (Huang et al. 2022). Out-of-the-box, ZairaChem models demonstrated state-of-the-art (SOTA) performance across all classification tasks, scoring between 1st and 7th in all benchmark datasets (Extended Data Table 2).

Figure 1 exemplifies ZairaChem applied to the H37Rv strain of *Mtb.* As of November 2021, 3,244 molecules had been screened in this assay at H3D, spanning a diverse chemical space (Fig. 1c). In total, 81 chemical series are represented, with 20 series covering 80% of the molecules. Setting an MIC_90_ cut-off of 5 μM yields 483 actives available for training the AI/ML model (Fig. 1b). To evaluate predictive performance, we held out 10% of the data as a test set and trained a model on the remaining 90%. An ensemble of models was fitted based on the multiple small-molecule descriptors and performance was evaluated for each model individually (Fig. 1f). The outcome of models inside the ensemble was aggregated in a consensus score estimating the probability of observing an “active” (1) assay outcome. Indeed, in the hold-out test set, known active molecules scored higher than known inactive compounds (Fig. 1d). Measured in the receiver operating characteristic (ROC) space, which evaluates the balance between sensitivity and specificity of a model, the consensus score achieves an area under the ROC curve (AUROC) of 0.895, higher than any of the individual classifiers alone. In this case, a score threshold of 0.25 can be established to classify predictions as “actives” (1) and “inactives” (0). Based on this threshold, the model predicted 62 active molecules from the test set, of which 37 were true positives, corresponding to a precision of 59.7% and a recall of 68.5% (Fig. 1h).

### Systematic AI/ML modelling of H3D screening cascades

We applied the ZairaChem pipeline to the experimental assays available at the H3D Centre (Fig. 2a and Extended Data Table 1). The resulting models showed good performance (AUROC > 0.7, Fig. 2b) and well-scaled prediction scores within the [0-1] range for all in-house datasets: *Pf* NF54, *Pf* K1, *Mtb* H37Rv, CHO, HepG2, Aqueous solubility (Aq. sol.), Caco-2 permeability (Caco-2) and Intrinsic clearance (CL_int_) for human (H), mouse (M), and rat (R) microsomes (Fig. 2c, Extended Data Fig. 1 and Table 3). We observed that ZairaChem classifiers successfully up-rank active compounds (Fig. 2d), with significant enrichment of hits within the top 50 candidates (Fig. 2e). We found similar performance for the remaining assays developed with in-house data beyond those depicted in Figure 2 (Extended Data Fig. 1, 2).

Data points were scarce for key assays related to more advanced stages of the screening cascade. Experiments related to drug metabolism, such as interactions with cytochrome P450 enzymes (CYPs), or inhibition of the hERG ion channel, are costly and often not performed on-site for many drug discovery organisations. In the case of CYPs, we gathered bioactivity data for over 15,000 molecules available from the PubChem BioAssay (Kim et al. 2022) and ChEMBL databases (Davies et al. 2015). We built AI/ML models for the CYP3A4, CYP2C9, CYP2C19 and CYP2D6 isoforms. Remarkably, except for CYP2C19, CYP models built with public data were able to achieve good performance (AUROC > 0.65) in the H3D chemical space (Fig. 2b and 2d, Extended Data Fig. 2, Table 3), assigning high scores to active (“1”, CYP inhibitors) and low scores to inactive compounds (“0”, no CYP inhibition observed) (Fig. 2c and Extended Data Fig. 1). These models successfully enable the selection of compounds that are not likely to interact with the selected CYPs (50 bottom-ranking compounds; Fig. 2e, Extended Data Fig. 3 and Table 4).

In addition to publicly available datasets, pre-trained AI/ML models are becoming more frequent in the scientific literature. The core mission of EOSI is to collect such public models in a unified, easy-to-use repository named the Ersilia Model Hub (Turon and Duran-Frigola, 2022). As of December 2022, the Ersilia Model Hub contains over eighty models for drug discovery, with a focus on infectious disease research. To demonstrate the potential of this resource, we chose to fetch a hERG blockade prediction model from the Ersilia Model Hub (identifier: eos2ta5), corresponding to CardioToxNet (Karim et al. 2021). This model was used “as is”, without further fine-tuning with H3D data, and showed excellent accuracy (AUROC = 0.852) on H3D compounds (Fig. 2b and d, Extended Data Table 3), with good discriminative scores between active (1, cardiotoxic) and inactive compounds (0, non-cardiotoxic) (Fig. 2c), as well as an enrichment of inactivity in the bottom-ranked 50 compounds (Fig. 2e).

Finally, to measure the advantage of using AI/ML models to prioritise compounds for experimental screening, we have calculated the hit enrichment potential of each model. Overall, assays where an “active” outcome is desired (*Pf* NF54, *Pf* K1, *Mtb* H37Rv, Aq Sol and Caco-2) show between 25% and 41% enrichment in the top 50 molecules. For example, out of 50 randomly-selected molecules in the *Pf* NF54 dataset, by chance, 10 would be active (20%); however, if we rank the data according to the AI/ML model scores, 42 molecules (84%) are found to be active (IC_50_ <0.1 μM). Assays where the desired outcome is “inactivity” (cytotoxicity, intrinsic microsomal clearance, CYP inhibition and hERG cardiotoxicity) show an enrichment of 30% to 50% of inactive molecules in the bottom 50 molecules ranked by the AI/ML model score (Extended Data Table 3, Extended Data Fig. 3 and 4). Moreover, we demonstrate how models trained on external datasets (P450 CYPs models) can improve their performance when in-house data points are included in the training set. Notably, the AUROC of the worst-performing model on H3D data (CYP2C19) goes from 0.54 to 0.7 upon adding internal data points (Extended Data Fig. 5).

### AI/ML performance by chemical series

In the early stages of a drug discovery project, derivatives of a compound which retain the structural core, or pharmacophore, are designed and synthesised to optimise the biochemical properties of that chemical series. Given the high performance of ZairaChem models for datasets spanning broad chemical space, we turned our attention towards the capability of the models to capture granular detail within a set of closely-related analogues, focusing on predictions of bioactivity against *Pf* and *Mtb.* We trained sets of models on bioactivity data and gradually increased the number of training points from specific chemical series to observe the effect of increasing “local” training data on model performance. As expected, the predictive potential for a chemical series improved with an increase in local data density, where approximately 30 molecules from a given series provide a good starting point to produce predictive models (Fig. 3c,g). Secondly, we measured the impact of the availability of global data on model quality at the chemical series level (Fig. 3d,h). In general, the addition of more data, even if they correspond to a broader chemical space, improves model performance. Series 2 of *Mtb*, the exception to this trend, had very few active compounds available for training, leading to poor model performance that the addition of global data could not rescue.

**Figure 3.**
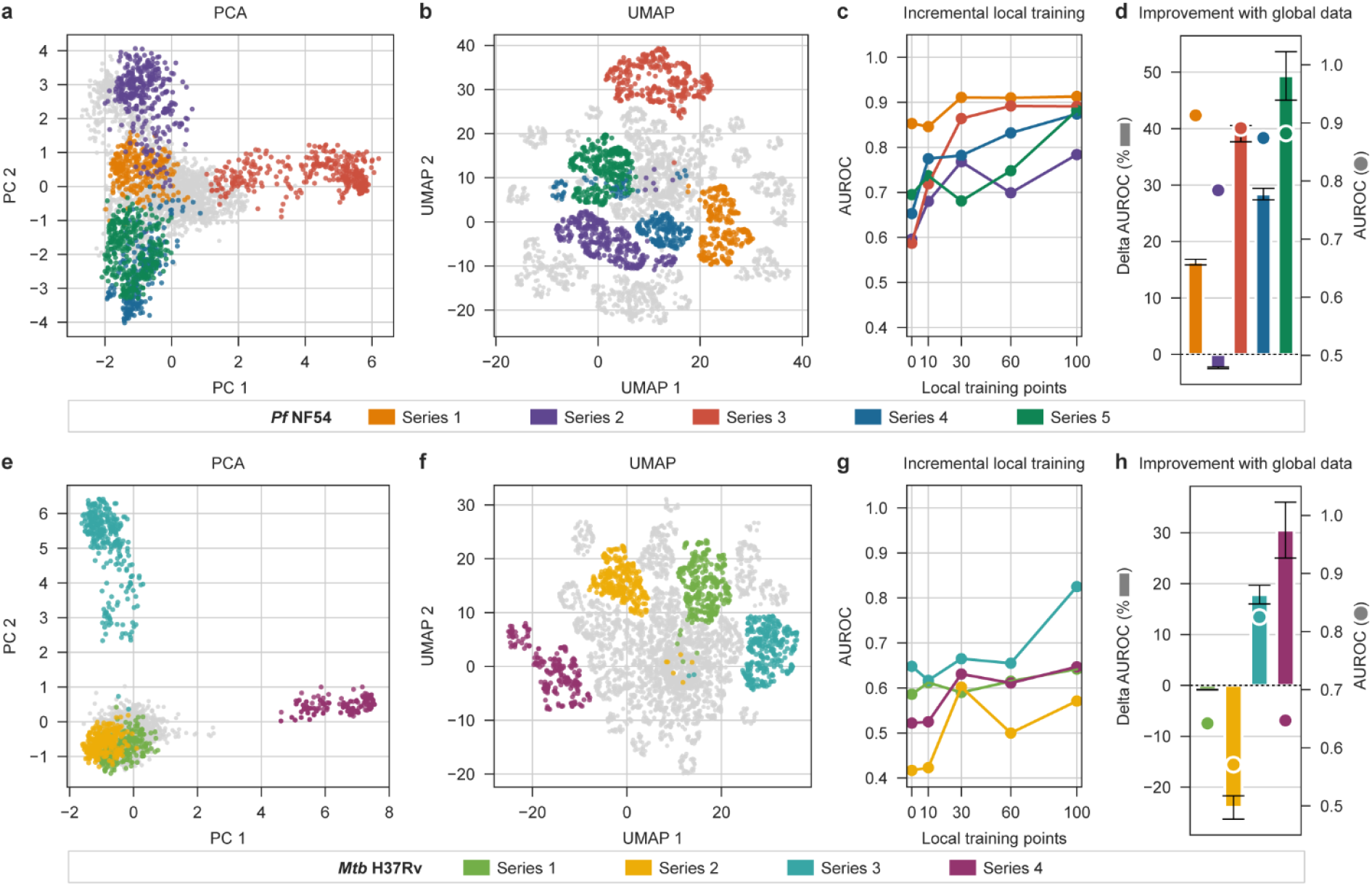
Model performance within chemical series corresponding to novel regions of chemical space. PCA and UMAP projections of the chemical space for the H3D Centre’s library depict compound distribution for specific chemical series in the malaria (top row) and tuberculosis (bottom row) disease areas: (**a** & **e**) PCA preserves the global distribution of chemical space while (**b** & **f**) UMAP emphasises the clustering of similar data points. (**c** & **g**) Median AUROC scores from a five-fold cross-validation are measured for training sets with an increasing number of local training points for each series, respectively. (**d** & **h**) The percentage of change towards a perfect model (AUROC = 1) between a model trained on a dataset that includes compounds from a more general chemical space versus a model trained on series-specific data alone (see calculation in Materials and Methods). The median AUROC score from a five-fold cross validation, for models trained with 100 series-specific compounds and global data, is plotted with a circle corresponding to the values of the right-hand-side y-axis.

### Application to an unseen collection of compounds

Finally, to demonstrate the effectiveness of the virtual screening cascade for *de novo* screening of molecular libraries, we reproduced the discovery of a potent antiplasmodial compound at H3D from a series of 2,4-disubstituted imidazopyridines (Horatscheck et al. 2020). In this study, the authors identified two initial hits with moderate antiplasmodial activity against asexual blood stage parasites (IC_50_ = 0.24 μM and 0.49 μM, respectively) and derived a series of 65 compounds for experimental testing. All molecules were tested against asexual blood stage drug-sensitive (NF54) and drug-resistant (K1) *Pf* strains, and their cytotoxicity in CHO cells as well as aqueous solubility (pH 6.5) were experimentally determined. In addition, some molecules progressed further in the cascade and were tested for cardiotoxicity risk (hERG blockade) and microsomal metabolic stability (Mouse CL_int_). To simulate a prospective study for this series, we removed these compounds from the training sets of the relevant AI/ML models in the virtual screening cascade (*Pf* NF54, *Pf* K1, CHO, Aq. Sol. at pH 6.5, Caco-2, CL_int_ Human, Rat, and Mouse). CYP and hERG models were developed with external data and thus do not contain any H3D molecules in the training set. ZairaChem models show good performance (AUROC > 0.75) on the experimentally validated library (Fig. 4a). Next, we leveraged the model scores to create visual fingerprints for ease of identification of potential hits: those with high scores for the desired activities (dark red) and low scores for the undesired activities (light blue) (Fig. 4b). We selected a few molecules that showed the desired pattern (Fig. 4c) and compared their predicted activity from the ZairaChem models with the experimental results (Fig. 4d). Compound 1, the initial hit of the series, is also shown for reference. We demonstrate that AI/ML models allow for the selection of candidates with high chances of progression in the cascade (compounds 37 and 55) and, conversely, molecules with undesired side effects could be discarded prior to experimental testing, despite their high bioactivity against *Pf* (compounds 22 and 58). Indeed, compound 37, one of the molecules selected by our model predictions for its high activity against *Pf* and low toxicity profile, was the lead compound from this study, showing *in vivo* efficacy in the humanised SCID NSG mouse malaria infection model.

**Figure 4.**
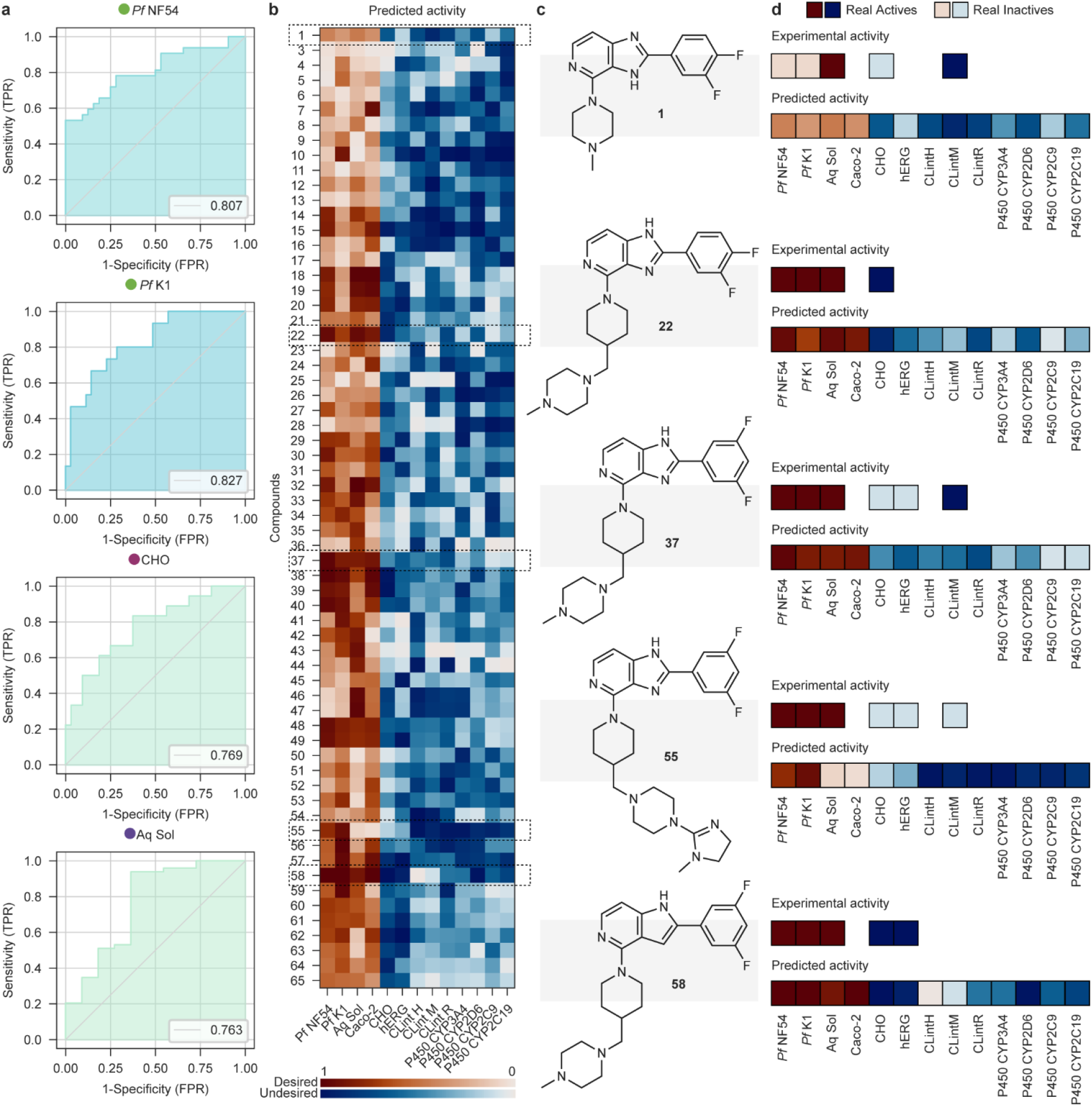
*De novo* screening of libraries using AI/ML models. (**a**) ROC curves of ZairaChem models tested on the library of 65 compounds (not included in the training set). Legend indicates the AUROC values of each model. Only models for which experimental validation was available for the 65 molecules are shown. (**b**) Predicted scores for each compound, transformed to a scale of 0 to 1 for comparison between assays. Desired activities are shown in a red colour scale and undesired activities are shown in a blue colour scale. Colour maps fade from 1 to 0 according to each model score. (**c**) Structure of selected compounds, including the initial hit compound 1. (**d**) Comparison of the predicted score and the real experimental activity of selected compounds (non-existing squares indicate no experimental data on these assays).

Experimental activity is represented as 1 (dark blue or dark red) or 0 (light blue, light red) for desired and undesired assay outcomes, respectively.

## Discussion

We have introduced ZairaChem, a robust and parameter-free AutoML tool that is broadly applicable to small-molecule property prediction tasks. ZairaChem makes use of a range of chemical descriptors in combination with an ensemble of AI/ML algorithms to train models with SOTA performance as an out-of-the-box tool. We implemented the tool systematically to the existing drug discovery pipeline at H3D, yielding 15 production-ready models corresponding to key assays related to antimalarial and antitubercular screening cascades. ZairaChem models, whether trained on in-house data or from public data sources, showed excellent performance, with most AUROC scores above 0.75. The resulting hit enrichment reduces attrition rates of the experimental pipeline, potentially accelerating the bench-to-bedside turnaround time. For example, here we show that by testing just 50 compounds of the whole-cell screening data, we find four times more active compounds against *Pf* (NF54 strain) and *Mtb* (H37Rv strain) compared to testing the same number of compounds selected at random. This is particularly relevant in the context of research conducted within LMICs, where resources are typically constrained.

Subsequent analysis of local (series-specific) regions of the chemical space aimed to answer two questions; firstly, whether our AI/ML models were able to capture differences between closely-related compounds within a specific chemical series and, secondly, whether the presence of broader “global” data contributed to the improvement of predictions in these series. With no local training data, the prediction quality of ranking analogues varies from series to series; some chemical series are readily modellable (*Pf* NF54 Series 1, Fig. 3c) while others require additional “local” training data to improve predictions (*Pf* NF54 Series 2, and 4, Fig. 3c). While a limited local training set of just ten compounds has the potential to skew model performance depending on the specific split of data points, having approximately 30 local molecules within a chemical series appears to be a useful rule-of-thumb to produce models that can distinguish between small variations in chemical structure. Additional data, whether local (Fig. 3c & 3g) or global (Fig. 3d & 3h), generally further enhances model performance.

Finally, we have demonstrated the potential of AI/ML virtual screening cascades to identify hits in a *de novo* library by recreating a study performed at the H3D Centre in 2020. The visual representation of compounds’ colour-vectors, based on AI/ML model scores, allowed for the quick identification and selection of the best compounds (highly potent against *Pf* with low toxicity) for experimental testing.

Overall, we have successfully deployed 15 AI/ML models at H3D. These models are intended to serve two main purposes. On the one hand, they are used to screen large compound libraries to discover new chemical matter as starting points for drug discovery projects. On the other hand, they provide a means to prioritise compounds for synthesis in the hit-to-lead and lead optimisation phases of existing screening campaigns. In the latter scenario, medicinal chemists frequently design dozens of potential compounds but have limited resources (including time) for chemical synthesis. The ZairaChem virtual screening cascade is intended to be used in conjunction with other existing software tools to generate complementary data points that facilitate decision-making, reduce attrition rates and accelerate project progression. As models can provide predictions for many molecules, scientists are able to explore much longer lists of compounds than they might otherwise propose, allowing them to interrogate a broader chemical space. Furthermore, predictions for design ideas can be obtained from the entire virtual screening cascade to identify compounds with potential benefits and liabilities much earlier in the design-make-test cycle, including those with predicted favourable ADMET assay outcomes that might typically only be considered later in a drug discovery programme.

Currently, ZairaChem is limited to single-output binary classification tasks. Although the framework is prepared to deal with regression (i.e. continuous data) tasks, we chose to restrict the scope to the former at this first stage of development. This choice matches decision-making guidelines at H3D, where expert cut-offs are defined for each assay to determine progression criteria. Likewise, although ZairaChem is now being extended in new directions, including interpretability, data augmentation for low-data regimes, and confidence estimation, these functionalities were not implemented in the current study, which is primarily focused on overall assessment of hit rates and predictive performance. Finally, ZairaChem can natively incorporate any AI/ML model available from the Ersilia Model Hub (Turon and Duran-Frigola 2022), including modern descriptors, embeddings extracted from pre-trained models such as GEM (geometry-enhanced molecular representation) (Fang et al. 2022), or models fitted to a broad range of bioactivities using, for instance, the FS-Mol training set (Stanley et al. 2021). Here, we chose four representative descriptors corresponding to physicochemical properties, classical 2D structure fingerprints, advanced graph embeddings and bioactivity signatures. Improvement or extension of this default set is outside the scope of the study; however, this could serve to further improve the performance of the models presented here. It is worth noting that, in addition, the Ersilia Model Hub contains a growing number of predictive models for specific assay outcomes, developed either by third parties or by EOSI using publicly-available data (e.g. from ChEMBL (Gaulton et al. 2017) or PubChem BioAssay (Kim et al. 2022)). For example, the MAIP antimalarial model (Bosc et al. 2021) is readily accessible, as well as popular ADMET servers (SwissADME, ADMETlab 2.0) (Daina, Michielin, and Zoete 2017; Xiong et al. 2021) and hERG inhibition models. ZairaChem can treat the outcome of these models as additional features for prediction. MAIP, for instance, can predict H3D *Pf* NF54 outcomes with AUROC = 0.73 as a standalone estimator. Thus, the ZairaChem framework offers a unique opportunity to transfer knowledge from the public domain to in-house modelling efforts.

## Concluding remarks

In summary, we have successfully developed and deployed an AI/ML-based virtual screening pipeline based completely on open-source code and conventional computing capacity. This is the first instance of a virtual screening cascade built with data produced on and for the African continent. We hope that the work presented here serves as a proof-of-concept for the potential of AI/ML tools to support drug discovery efforts in LMICs. Incipient centres in West Africa (Amewu et al. 2022) and Central Africa (Namba-Nzanguim et al. 2022) may benefit from similar implementations. More globally, ZairaChem offers a competitive, free and constantly-updated software solution to model small-molecule bioactivity data, where no strong data science skills are required to run the tool.

In order to encourage scientists at H3D to use our models, a series of internal and external webinars have been held to introduce AI/ML in the context of drug discovery and to explain how the models were trained as well as how to interpret the results (see, for example, the recent “Bringing data science and AI/ML tools to infectious disease research” workshop co-organized by the H3D Foundation and EOSI; https://ersilia.gitbook.io/event-fund). “Champions” in each core team at H3D helped to promote the tools and brought ideas and suggestions back to the development team. In this rollout phase of the implementation, a list of compounds was sent to the AI/ML team who employed the pipeline and then shared the results with the medicinal chemistry project team. In the longer term, it is envisaged that scientists will be able to run predictions on their own. We also anticipate expanding the array of available models and building models on demand, especially for target-based projects such as those based on the clinically-validated *Pf* phosphatidylinositol 4-kinase (PI4K) enzyme (Arendse et al. 2022; Paquet et al. 2017). Continued revision and maintenance of the models ensure that they are updated with the latest data being generated at H3D. Likewise, contrary to many ‘static’ AI/ML tools published in an academic setting, ZairaChem is in continuous development, including novel descriptor types and algorithms over time.

Finally, plans for the future include the coupling of ZairaChem AI/ML assets to generative models, with the goal to design compounds *de novo* and/or aid the hit-to-lead optimisation step. In the context of the Open Source Malaria (OSM) and Open Source Antibiotics (OSA) consortia, EOSI has demonstrated how a “reinforcement learning” scheme can be applied to generate compounds with predicted desired biochemical properties^2^. Thus, we envisage a wet-lab/dry-lab cycle implemented at H3D whereby AI/ML models are trained based on historical in-house data and then incorporated into a generative framework to suggest new small-molecule candidates with optimised prediction outputs. Upon clerical inspection by medicinal chemists, candidates would eventually be synthesised and tested. Results of the assay will in turn be incorporated in the next training round, improving the AI/ML model and its correspondence with the existing experimental screening facilities. The current successful and sustained implementation of ZairaChem at H3D represents the first step towards truly integrated AI/ML usage in productive drug discovery settings in the Global South.

## Materials and Methods

### Data collection

Bioactivity data for the assays defined in Extended Data Table 1 were extracted from H3D’s curated database. Compounds with large variations in replicated measurements were identified and removed. The replicate values for remaining compounds were averaged regardless of date and site of experiment development.

Cytochrome inhibition data was obtained for CYP3A4, CYP2C9, CYP2C19, and CYP2D6 from PubChem (AID1851, AID899, AID891, AID883, AID884) and ChEMBL. PubChem BioAssay data were already binarized (1, active; 0, inactive). The ChEMBL database was queried and only compounds with “Standard Type”, “IC_50_” or “*K*_i_”, and “Standard Units”: “nM” were considered. Compounds were assigned as “active” (1) if bioactivity was smaller or equal to 10 μM, and “inactive” (0) otherwise. In addition, compounds with “Comment” equal to “Not Active” or “No Inhibition” were classified as inactive. Bioactivity data not matching these criteria were discarded.

### The ZairaChem pipeline

Each H3D assay was modelled independently using ZairaChem with default parameters. ZairaChem has two running modes, namely “fit” and “predict”. ZairaChem training runs are scheduled to be executed twice a year for all assays in the virtual screening cascade.

#### Data pre-processing

The data pre-processing module consists of several steps. First, an input file is analysed and relevant columns are identified. In particular, the column containing SMILES strings is kept, together with the outcome (e.g. activity) column. ZairaChem determines the type of task (i.e. regression or binary classification). In this study, only binary classification tasks were considered. Small-molecule SMILES are standardised following the MELLODDY-Tuner protocol (Oldenhof et al. 2022). MELLODDY-Tuner is also used to identify LSH-based as well as Murcko scaffold-based splits, which can be used optionally. In addition, ZairaChem enables random and time-based splits, as well as a splitting scheme that takes into account clusters and LSH hashes to identify equally-sized splits. By default, five random stratified splits are done, keeping 10% of the data as test sets.

#### Small-molecule descriptors

ZairaChem can query the Ersilia Model Hub, EOSI’s repository of pre-trained, ready-to-use AI/ML models. Some of the assets available in the Ersilia Model Hub provide as output a numerical vector for each molecule, typically capturing physicochemical or topological characteristics of the compound. By default, ZairaChem calculates such vectors for each molecule, corresponding to four representative descriptor types. In particular, we calculate (1) Mordred descriptors (an array of >1600 physicochemical parameters), (2) ECFP fingerprints (a count-based vector of 2048 dimensions based on circular exploration (radius 3) of all atoms in a molecule), (3) Chemical Checker signatures (a dense vector of 3200 dimensions capturing known and inferred bioactivity data across a wide range of bioactivity outcomes), and (4) GROVER embeddings (a 5000-dimension graph-based representation of the molecules). Other vectorial descriptors are available in the Ersilia Model Hub and can be specified to ZairaChem by simply referring to their model identifier.

Continuous data descriptors are quantile-normalised and missing data is imputed with a nearest-neighbour approach. Invariant columns are removed. GROVER is used as a reference descriptor for additional processing. We perform PCA (four components) and UMAP (two components). In addition, supervised versions of these techniques are applied based on the binary outcomes. We choose linear optimal low-rank projection (lolP) (Vogelstein et al. 2021) as an alternative to PCA for this supervised task. UMAP accepts both unsupervised and supervised modes. In addition to vector-like descriptors, it is possible to incorporate other models from the Ersilia Model Hub as auxiliary predictor variables for ZairaChem. ZairaChem treats these auxiliary models as additional columns in the pooling step described below.

#### AutoML methods

Currently, ZairaChem executes four AutoML methods independently. Each of the AutoML models is focused on enhancing a specific feature (e.g. interpretability, robustness, etc.) in the overall ZairaChem pipeline.

The first AutoML module performs independent modelling for each of the descriptors. The focus of this module is to identify which descriptor types are the most appropriate for the task of interest. With default parameters, five models are built, corresponding to the descriptors mentioned above. FLAML (Wang et al. 2019) is used to perform rapid search of Random Forest hyperparameters.

The second module is focused on visual interpretation of the chemical space. Thus, this module takes as input the low-dimensional PCA and UMAP projections obtained for the reference descriptor (in total, 12 variables). We use AutoGluon-Tabular (Erickson et al. 2020) with default parameters to obtain robust classifiers and regressors.

The third module leverages GROVER, a data-driven descriptor trained on a large collection of molecules. This module is illustrative of a “transfer learning” approach where a large chunk of a neural network is “frozen” (the GROVER part) and a few extra layers are fine-tuned for the task of interest. Here, we add an additional dense layer. The number of dimensions of this layer, in addition to the training parameters, are automatically selected with Keras Tuner.

Finally, the fourth module leverages image-based representations of the molecules, enabling application of computer vision techniques, which are particularly advanced in the field of AI/ML. Compounds are represented as MolMaps (Shen et al. 2021). These are concise, multi-descriptor images (maps) where regions of the image correspond to descriptors that are correlated. For example, in a MolMap one can find a region that relates to size and molecular weight, whereas other regions are related to lipophilicity, solubility, etc. It has been shown that convolutional neural networks can be used out-of-the-box taking MolMaps as input, without the need for intense architecture and hyperparameter search.

ZairaChem is prepared to be extended with additional AutoML modules, if and as necessary. However, the balance between computing time and gain in performance is an important consideration before adding further modules.

#### Pooling

Each of the models above provides point predictions that can be aggregated in a consensus (pooled) prediction. By default, ZairaChem applies a “blending” approach, based on a weighted average between individual predictions.

#### Reports and output

At the end of the ZairaChem pipeline, performance reports are automatically provided, including the most common validation metrics for binary classification tasks. A single spreadsheet with prediction output and performance metrics is provided as a primary result, along with multiple reporting plots (AUROC, UMAP projections, etc.).

#### ZairaChem benchmarking

We selected Therapeutics Data Commons (TDC) as a benchmark framework. TDC contains multiple datasets across a broad range of tasks and activities. To prove SOTA performance on the H3D Centre’s data, we selected the TDC ADMET Group as the one containing tasks most similar to the ones described in this project. In line with this, only classification tasks were evaluated. Data was downloaded from TDC and split into train and test according to the TDC guidelines. Models were trained with default ZairaChem parameters with 8-fold cross validation. Results were calculated using TDC guidelines.

### Analysis of H3D’s chemical series

To investigate the change in model performance within a localised region of chemical space through the incremental addition of local training data, chemical series with at least 200 compounds were selected from the H3D library for analysis.

#### Train-test splits

Datasets for model training were constructed according to the following protocol: first, all compounds in a series of interest were removed from the bulk global training data of the H3D library; then, the data was shuffled and 100 compounds were randomly selected with stratification as a standard test set; the remaining 100 molecules were systematically added back to the bulk H3D activity data and models trained for each dataset. A final separate ‘local-only’ model was trained on the 100 series-specific compounds alone, without the broader H3D library data, in order to investigate the effect of global training data on model performance.

#### AUROC percentage change

To measure the contribution of additional global training data to a model’s performance in a localised chemical space, we calculate the percentage change in AUROC score according to the following steps: 1) first, we find the difference between the AUROC scores of the model trained on 100 series-specific compounds with the H3D library included as well as the ‘local-only’ model trained on the series-specific compounds alone; 2) next, for the ‘local-only’ model, we find the AUROC score that is still possible to be achieved (1 - ‘local-only’ AUROC); 3) lastly, we take the difference in models scores (from step1) as a percentage of the score that could still be achieved (from step 2). This metric represents the additional performance gained or lost out of the total AUROC score that was still available to capture. AUROC scores were calculated with the SciKit-Learn package from a five-fold cross-validation.

#### Plots

Chemical space visualisation was constructed by describing compounds using the Morgan Fingerprint algorithm, with a radius of 3 and vector length of 2048 as implemented in the RDKit package. These fingerprints were then projected onto two dimensions through principal component analysis as well as the UMAP algorithm and plotted with the Matplotlib library.

## Data and code availability

ZairaChem is available at https://github.com/ersilia-os/zaira-chem. A command-line interface to the Ersilia Model Hub is accessible at https://github.com/ersilia-os/ersilia, and the repository can be browsed from https://ersilia.io/model-hub. Extended documentation can be found in the Ersilia Book (https://ersilia.gitbook.io). ZairaChem benchmarks can be found in https://ersilia-os/zaira-chem-tdc-benchmark. Figures were styled using EOSI’s Stylia library (https://github.com/ersilia-os/stylia). Code used for analysing data can be found at https://github.com/ersilia-os/h3d-screening-cascade-code

## Author contributions

MDF, GT and KC designed the study. MDF developed ZairaChem with the support of JH and GT. GT performed the benchmarking. GT curated and analysed the data with the help of JW and JH. MDF, GT and JH performed the analysis. JH and JW implement and maintain the models at H3D. KC and MDF supervised the project. All authors discussed the results and commented on the manuscript.

## Acknowledgements

EOSI is grateful to Merck KGaA for a Biopharma Speed Grant. JH is a recipient of the Harry Crossley Foundation Postdoctoral Fellowship. Capacity building activities within the context of the H3D-EOSI collaboration were supported by an Event Fund grant from Code for Science & Society and the Wellcome Trust. We thank Dr Susan Winks for her support in coordinating this event. KC is the Neville Isdell Chair in African-centric Drug Discovery and Development and thanks Neville Isdell for generously funding the Chair. The South African Medical Research Council and South African Research Chairs Initiative of the Department of Science and Innovation are gratefully acknowledged for their support (KC). The authors would like to thank Dr Jake M. Pry for providing feedback on the manuscript, Dr André Horatscheck and Dr Grant Boyle for kindly facilitating access to the central database at H3D, as well as Dr Preshendren Govender for sharing the *M. tuberculosis* screening data.

## Footnotes

1 Both Ersilia and Zaira are cities described in Italo Calvino’s book *Invisible Cities* (1972). Ersilia is a “trading city” where inhabitants stretch strings from the corners of the houses to establish the relationships that sustain the life of the city. When the strings become too numerous, they rebuild Ersilia elsewhere, and their network of relationships remains. Zaira is a “city of memories”. It contains its own past written in every corner, scratched in every pole, window and bannister.

2 https://github.com/ersilia-os/osm-series4-candidates-round-2

