## Extended_Data for "First fully-automated AI/ML virtual screening cascade implemented at a drug discovery centre in Africa"

| Assay | Description | N. molecules | Activity cut-off |
|---|---|---|---|
| *P. falciparum* NF54 $IC_{50}$ | Half-maximal inhibitory concentration of *P. falciparum* NF54 strain cultured at 5% parasitemia in human blood | 3289 | 0.1 μM |
| *P. falciparum* K1 $IC_{50}$ | Half-maximal inhibitory concentration of *P. falciparum* K1 strain cultured at 5% parasitemia in human blood | 1425 | 0.1 μM |
| *M. tuberculosis* $MIC_{90}$ | Minimal inhibitory concentration 90% of *M. tuberculosis* H37Rv strain cultured in glucose-Tween80 media | 3244 | 5 μM |
| CHO $IC_{50}$ | Half-maximal inhibitory concentration of CHO cell growth | 2029 | 10 μM |
| HepG2 $IC_{50}$ | Half-maximal inhibitory concentration of HepG2 cell growth | 1457 | 10 μM |
| $CL_{int}$ Human | Human microsomal clearance *in vitro* | 1430 | 11.6 μg/min/mg |
| $CL_{int}$ Mouse | Human microsomal clearance *in vitro* | 1165 | 11.6 μg/min/mg |
| $CL_{int}$ Rat | Human microsomal clearance *in vitro* | 1202 | 11.6 μg/min/mg |
| Aqueous solubility | Solubility at pH 7.4 | 3227 | 90 μM |
| Caco-2 | Passive membrane permeability | 134 | 10 μM |
| CYP2C9 $IC_{50}$ | Half-maximal inhibitory concentration of cytochrome CYP2C9 (from literature) | 16379 | 10 μM |
| CYP2C19 $IC_{50}$ | Half-maximal inhibitory concentration of cytochrome CYP2C19 (from literature) | 15551 | 10 μM |
| CYP2D6 $IC_{50}$ | Half-maximal inhibitory concentration of cytochrome CYP2D6 (from literature) | 17812 | 10 μM |
| CYP3A4 $IC_{50}$ | Half-maximal inhibitory concentration of cytochrome CUP3A4 (from literature) | 21810 | 10 μM |
| hERG $IC_{50}$ | Blockade of the human ether-a-go-go-related gene potassium channel (from literature) | 12620 | 10 μM |

**Extended Data Table 1**. Bioassay descriptions with dataset sizes and activity cut-offs for the classification models.

| Dataset | Metric | Score | Position* |
|---|---|---|---|
| Bioavailability_Ma | AUROC | 0.706 ± 0.031 | 3rd |
| HIA_Hou | AUROC | 0.948 ± 0.018 | 7th |
| Pgp_Broccatelli | AUROC | 0.935 ± 0.006 | 2nd |
| BBB_Martins | AUROC | 0.91 ± 0.024 | 2nd |
| CYP2C9_Veith | AUROC | 0.786 ± 0.004 | 3rd |
| CYP2D6_Veith | AUROC | 0.644 ± 0.085 | 4th |
| CYP3A4_Veith | AUROC | 0.875 ± 0.002 | 3rd |
| CYP2C9_Substrate_CarbonMangels | AUROC | 0.441 ± 0.033 | 1st |
| CYP2D6_Substrate_CarbonMangels | AUROC | 0.685 ± 0.029 | 2nd |
| CYP3A4_Substrate_CarbonMangels | AUROC | 0.63 ± 0.008 | 6th |
| hERG | AUROC | 0.856 ± 0.009 | 3rd |
| AMES | AUROC | 0.871 ± 0.002 | 1st |
| DILI | AUROC | 0.925 ± 0.005 | 3rd |

**Extended Data Table 2.** ZairaChem model performance on the Therapeutics Data Commons ADMET Leaderboard. The score is the mean ± standard deviation of the indicated metric on 8-fold cross validation. *Position refers to expected position in each leaderboard at time of submission.

| Assay | AUROC | Standard deviation |
|---|---|---|
| *P. falciparum* NF54 $IC_{50}$ | 0.915 | 0.020 |
| *P. falciparum* K1 $IC_{50}$ | 0.853 | 0.022 |
| *M. tuberculosis* $MIC_{90}$ | 0.901 | 0.036 |
| CHO $IC_{50}$ | 0.845 | 0.032 |
| HepG2 $IC_{50}$ | 0.973 | 0.010 |
| $CL_{int}$ Human | 0.794 | 0.027 |
| $CL_{int}$ Mouse | 0.782 | 0.056 |
| $CL_{int}$ Rat | 0.790 | 0.033 |
| Aqueous solubility | 0.888 | 0.019 |
| Caco-2 | 0.951 | 0.042 |
| *CYP2C9 $IC_{50}$ | 0.761 | 0.035 |
| *CYP2C19 $IC_{50}$ | 0.542 | 0.035 |
| *CYP2D6 $IC_{50}$ | 0.681 | 0.035 |
| *CYP3A4 $IC_{50}$ | 0.715 | 0.070 |
| *hERG $IC_{50}$ | 0.852 | 0.014 |

**Extended Data Table 3.** Model performances (area under the ROC curve (AUROC) and standard deviation). Models have been evaluated on 10% of the total data and 5-fold cross-validated. * Models developed with external data and models from the literature have been validated on 75% of the internal data at each fold. All splits have been stratified by actives/inactives.

| Assay | Enrichment in active compounds in the top 50 (%) | Enrichment in inactive compounds in the bottom 50 (%) |
|---|---|---|
| *P. falciparum* NF54 $IC_{50}$ | 41.8 | 49.22 |
| *P. falciparum* K1 $IC_{50}$ | 23.78 | 47.24 |
| *M. tuberculosis* $MIC_{90}$ | 37.84 | 47.18 |
| CHO $IC_{50}$ | 22.84 | 49.18 |
| HepG2 $IC_{50}$ | 47.6 | 49.42 |
| $CL_{int}$ Human | 42.5 | 39.52 |
| $CL_{int}$ Mouse | 37.4 | 29.62 |
| $CL_{int}$ Rat | 40.42 | 35.6 |
| Aqueous solubility | 49.46 | 47.56 |
| Caco-2 | *insufficient number of test molecules | *insufficient number of test molecules |
| *CYP2C9 $IC_{50}$ | 10.82 | 43.2 |
| *CYP2C19 $IC_{50}$ | 12.76 | 36.26 |
| *CYP2D6 $IC_{50}$ | 9.84 | 42.18 |
| *CYP3A4 $IC_{50}$ | 9.84 | 44.18 |
| *hERG $IC_{50}$ | 45.36 | 39.66 |

**Extended Data Table 4.** Hit enrichments in the top 50 and bottom 50 molecules ranked according to the model score (probability of 1).

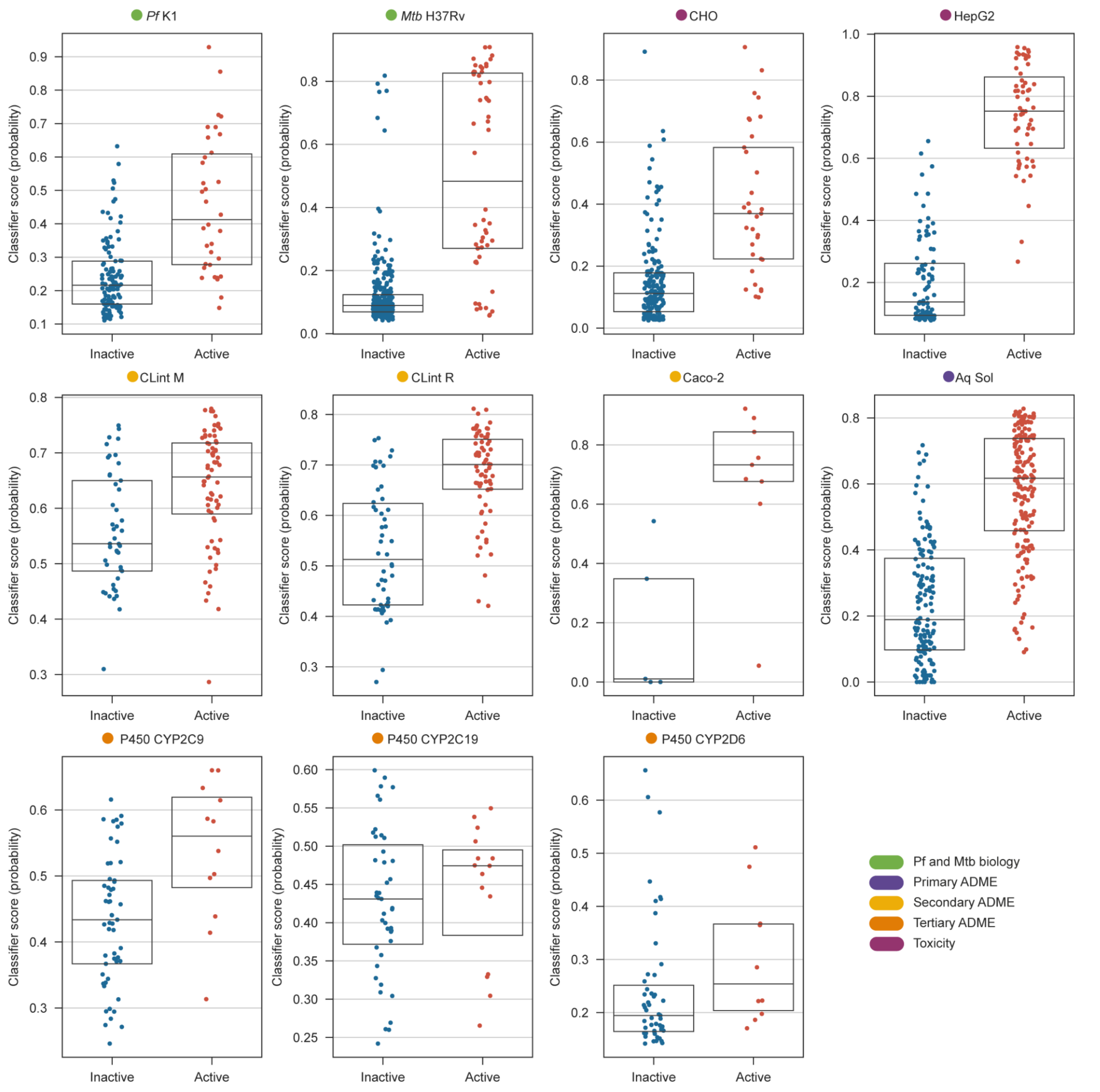


**Extended Data Figure 1.** Scores (probability of 1) obtained by ZairaChem classification models on test data (10% of total dataset). Blue indicates true inactive molecules and red represents true active molecules. Only one representative fold is shown.

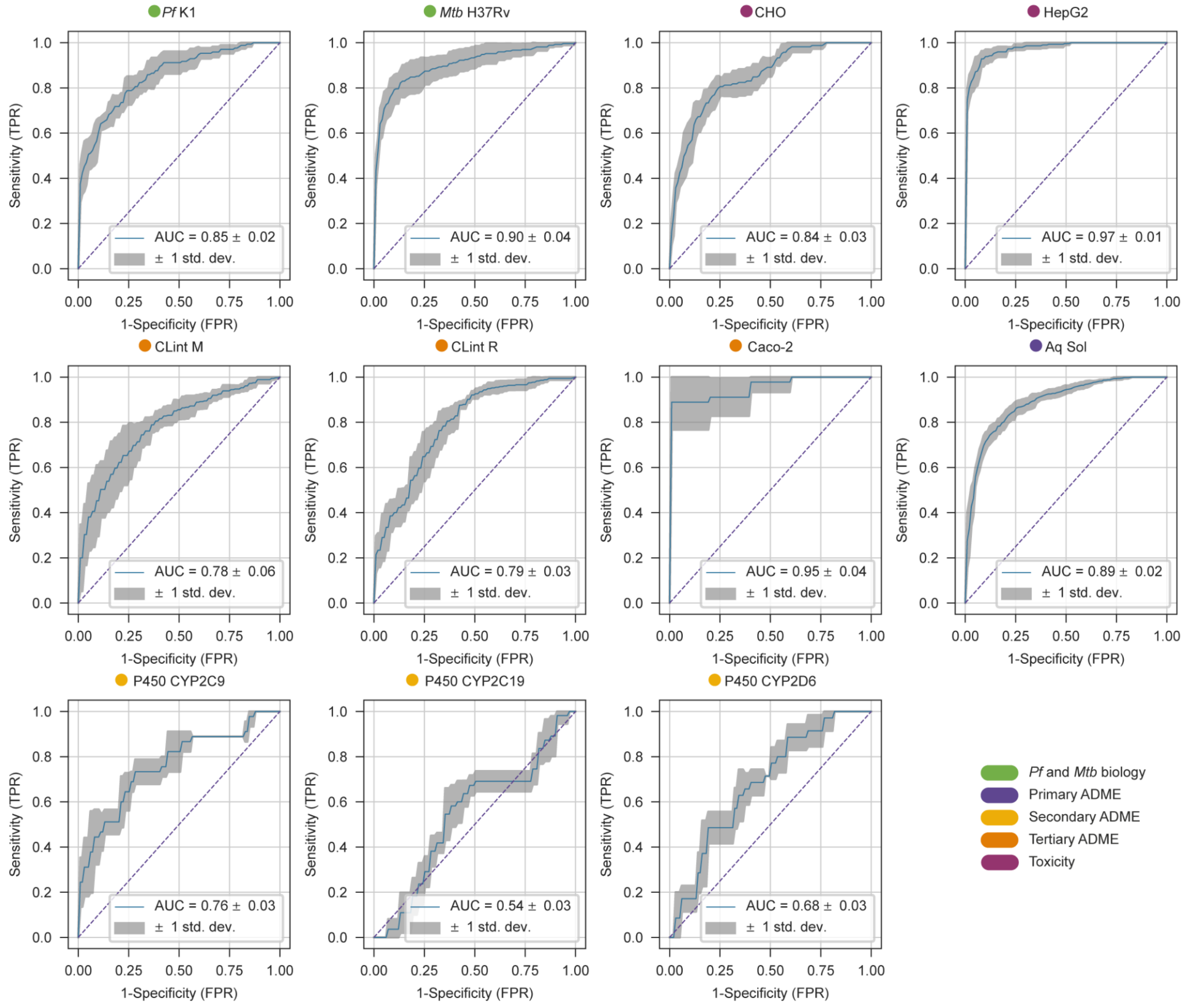


**Extended Data Figure 2.** ROC curves of 11 ZairaChem models and associated AUC values ± standard deviations (std. dev.). Models have been five-fold cross-validated using a random stratified 90-10 split. Blue lines represent the mean of the five folds.

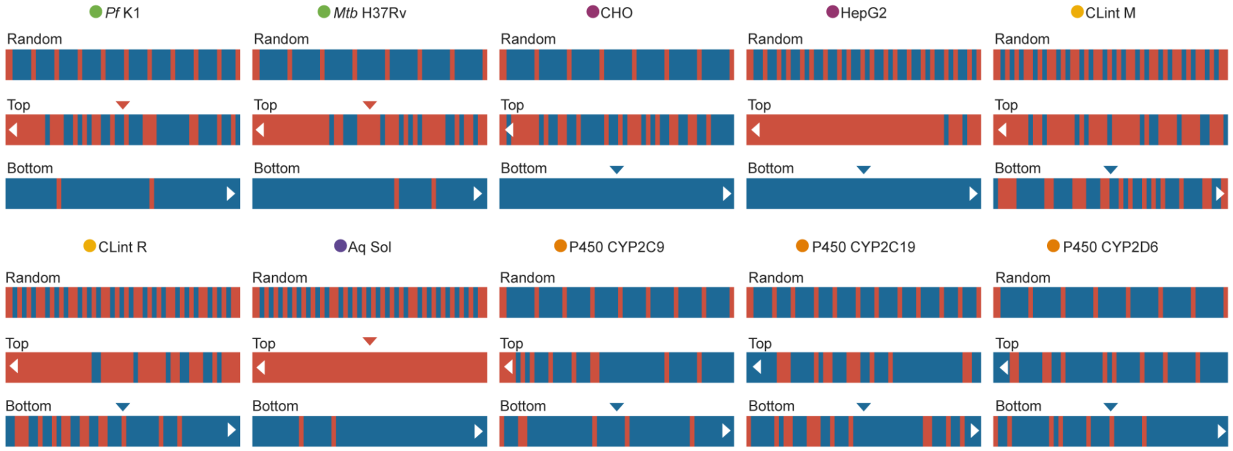


**Extended Data Figure 3.** Comparison of hit rates for randomly selected molecules (first row) vs molecules ranked according to the model score (probability of 1, second and third rows) for ten assays corresponding to activity against *Pf* and *Mtb*, cytotoxicity, intrinsic microsomal clearance in mice (M) and rats (R), aqueous solubility, and inhibition of CYP enzymes. The top 50 and bottom 50 molecules are depicted, showing a hit enrichment of true active compounds (red) in the highest-ranked positions and an enrichment of true inactive compounds (blue) in the lowest-ranked compounds. Red and blue arrows, respectively, represent the desired experimental outcome for molecule progression in the cascade.

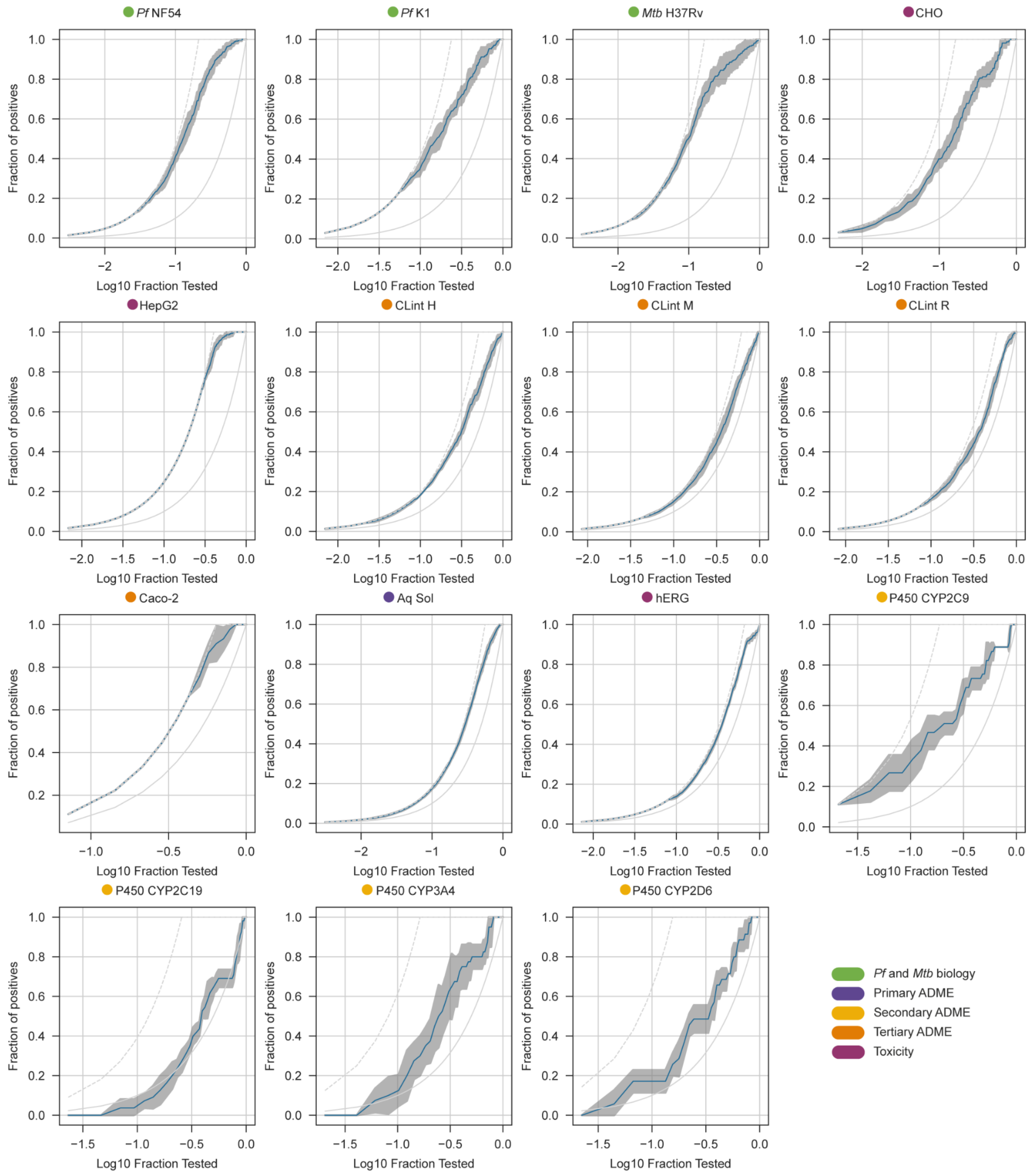


**Extended Data Figure 4.** Hit enrichment curves for the 15 models deployed as a virtual screening cascade. Blue lines represent the mean hit enrichment of the five-fold model cross-validations ± standard deviation (grey areas). Left and right grey lines represent the ideal situation (all actives are identified first) and the random situation (all molecules in the test set must be screened to identify all the active molecules), respectively.

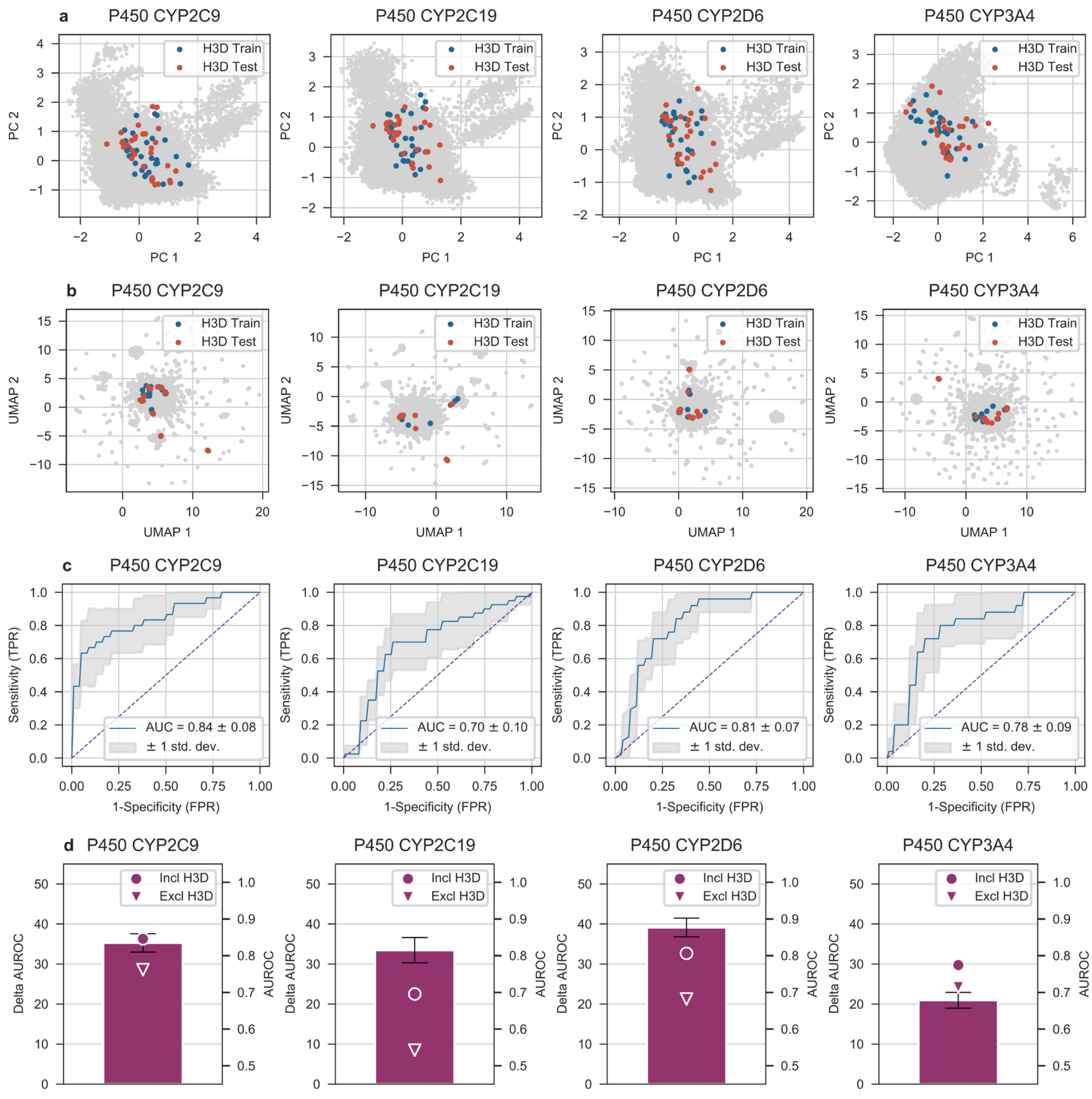


**Extended Data Figure 5.** Zairachem models for CYP P450 inhibition trained on external datasets, including 50% of internal H3D data points (approximately 30 molecules). The distribution of chemical space for each cytochrome dataset is depicted as two-dimensional projections with (**a**) Principal Component Analysis and (**b**) Uniform Manifold Approximation and Projection. These projections correspond to the first data fold from a five-fold cross validation, with public data (gray), H3D data included in the training set (blue) and H3D data used for the test set (red). (**c**) The mean ROC curves from the five-fold cross validation with standard deviations. (**d**) Percentage change in AUROC score (left y-axis) towards a perfect model (AUROC = 1) when adding internal data to the external training data (see ‘AUROC percentage change’ in Materials and Methods for analogous calculation). The right y-axis shows the actual AUROC values for the models before (downward triangle) and after (circle) adding internal data.
